# Telomere-to-telomere assembly detects genomic diversity in Canadian strains of *Borrelia burgdorferi*

**DOI:** 10.1101/2025.07.18.665644

**Authors:** Atia B. Amin, Ana Victoria Ibarra Meneses, Simon Gagnon, Georgi Merhi, Martin Olivier, Momar Ndao, Mathieu Blanchette, Christopher Fernandez-Prada, David Langlais

## Abstract

*Borrelia burgdorferi,* the bacteria causing Lyme disease, has a complex genome comprising a linear chromosome and a combination of linear and circular plasmids. The atypical hairpin structure at the telomere of linear replicons and the highly paralogous plasmids make the genome assembly challenging. We developed a genome assembly pipeline using Oxford Nanopore Technologies (ONT) long read and Illumina short read to overcome these challenges. Using ONT reads enabled us to completely assemble the hairpin telomeres of the linear replicons along with a novel lp28 subtype plasmid and the complete circular plasmids of nine *B. burgdorferi* strains from five geographical regions in Canada. Although these strains are highly similar across the conserved genomic regions, variability was observed predominantly at the right telomeric ends. Comparative analyses revealed that all nine strains carry a ∼2-10 kb right telomeric end identical to the linear plasmid lp28-1, which leads to variability in the telomere length and the gene content. Additionally, we observed diversity at the hairpin telomeric sequences of the linear chromosomes. Further analysis showed that the nine strains belong to seven *ospC* types and have diverse plasmid profile, highlighting the genomic diversity among the strains from the same geographical locations. Overall, these findings suggest that even the *B. burgdorferi* strains from close geographical locations can carry substantial genomic variation, especially at the telomeres and with respect to their plasmid content, emphasizing it to be a possible mechanism of rapid evolution within these Canadian strains.

## Introduction

Lyme disease (LD), caused by the spirochete bacteria *Borrelia burgdorferi* (*Bb*) sensu lato genospecies complex, is rapidly increasing in Canada due to climate change and the associated northward expansion of black-legged ticks, *Ixodes scapularis* (Bouchard et al. 2019; Russell et al. 2024; Kelly et al. 2024). Over the last three decades, LD has emerged as a major public health concern in Canada as the reported human LD cases rose from just 35 in 1994 to over three thousand cases in 2021 (Public Health Agency Canada, 2023), although actual case numbers are likely underestimated due to under-reporting and the challenges associated with diagnosing early-stage Lyme disease (Ogden et al. 2019). As a result, Lyme disease is now recognized as the most common vector-borne zoonosis in Canada, necessitating ongoing public health efforts to enhance surveillance, raise awareness, and develop effective prevention strategies (Wilson et al. 2022). While the complete genome sequences for endemic strains of *Bb* are important to develop protective measures, like diagnostic test to track the infection status and treatment success, these are critically limited for Canadian strains. To date, mostly partial *Bb* genomes are available for samples collected from Southern Manitoba, Northwest Ontario, and Nova Scotia (Tyler et al. 2018), leaving most parts of Canada unexplored. Although three draft genomes of *Bb* isolates from Western Canada have recently been published (Haidl et al. 2024), complete telomere-to-telomere assemblies and in-depth comparative analyses of the endemic Canadian strains are still lacking.

Assembling a complete genome of the genus *Borrelia* is a challenging task for several reasons. Although its genome is relatively small (∼1.5 Mb), it is highly fragmented and complex (Ren et al. 2023). Unlike any other bacteria, the *Borrelia* genome has a linear chromosome (size ranges from 900 to 920 kb), which has covalently closed hairpin loop ends as a special mechanism for solving the end-replication problem (Casjens et al. 1997; Tourand et al. 2009; Kobryn and Chaconas 2014). Interestingly, this type of hairpin telomeres is uncommon in prokaryotes, making *Borrelia* a powerful model organism to study hairpin end biology (Chaconas and Kobryn 2010). Unfortunately, our knowledge about the *Borrelia* hairpin telomere sequences across various *Borrelia* species and strains remained limited (Tourand et al. 2009). Most studies investigating the telomere sequences of *Borrelia burgdorferi* have predominantly utilized traditional methodologies, such as cloning following nuclease treatment; however, these approaches have limitations in confirming the precise nucleotide sequences at the hairpin turnaround, resulting in uncertainty regarding the nucleotide composition at the terminal ends (Huang 2004; Zhang et al. 1997; Tourand et al. 2009). Moreover, the hairpin loop wraparound is difficult to sequence using traditional short-read sequencing. Therefore, most of the assembly approaches using only Illumina reads could not assemble the linear genome up to the telomere due to low coverage at the hairpin loop. With the advancement of sequencing technology, we can now deploy long-read sequencing to solve this issue. (Hepner et al. 2023) already shown the advantage of using long PacBio reads to identify the telomeric inverse duplication in *Borrelia* genomes, whereas (Faith et al. 2024) resolved the hairpin telomeres of linear replicons in the *Bb* strain CA-11.2A genome using ONT long read.

Another unusual aspect of the *Borrelia* genome is its combination of more than 10 linear and circular plasmids, which together contribute to ∼40% of the genome (Margos et al. 2017). The plasmid composition varies between strains and plays a crucial role in virulence and adaptation to both the tick vector and mammalian hosts (Purser and Norris 2000). For example, some of these plasmids encode surface proteins, such as *ospC* and *vlsE*, which are essential for immune evasion and host interactions. Moreover, the linear plasmids show a wide range of variability in stability and some plasmids, such as lp28-1, lp25, and lp56, are easily lost during *in vitro* propagation (Margos et al. 2017). Similar to the linear chromosome, the linear plasmids also contain covalently closed hairpin telomeres (Barbour and Garon 1987; Casjens 1999), which adds complexity to *de novo* assembly.

A striking difference between the linear chromosome and linear plasmids is the degree of gene and sequence conservation. While linear chromosome is largely conserved and co-linear with the reference strain B31, linear plasmids show high variability in gene content and type across different species and strains (Casjens et al. 2012). This heterogeneity arises from the genetic rearrangements among plasmids, leading to a complex mosaic structure (Hepner et al. 2023). An example of such rearrangement is plasmid fusion, where a resulting plasmid carries identical sequences from multiple plasmid types. This high level of sequence homology, combined with the mosaic structure and similarity in plasmid sizes, presents significant challenges for *de novo* genome assembly (Casjens et al. 2000).

Due to this complexity, short-read sequencing alone is insufficient to assemble *Borrelia* plasmids (Margos et al. 2017; Becker et al. 2020; Hepner et al. 2023). A combination of long- and short-read sequencing could be more effective in this regard (Hepner et al. 2023), where the long reads can resolve the high level of homology across different plasmids by nearly spanning the entire plasmid as a single DNA sequence (Margos et al. 2017), while the short reads improves accuracy by polishing the assembled contigs. Beyond sequencing approaches, an efficient plasmid assembly pipeline is also important. Given that *Borrelia* carries numerous linear plasmids with terminal hairpin loops, assembly pipelines must be designed to accurately reconstruct these structures, while also reliably detecting terminal direct duplication in circular plasmid contigs.

In this study, we developed a hybrid genome assembly pipeline using Oxford Nanopore Technology (ONT) long reads and Illumina short reads to assemble nine *B. burgdorferi* isolates from five geographical locations across Northwest Ontario and Manitoba. Our goal was to assemble a set of complete genomes from these endemic regions, including the linear chromosome and the full complement of linear and circular plasmids. To achieve this, we developed a streamlined bioinformatics pipeline that is tailored for high-confidence assembly of *Borrelia*’s chromosome and replicons. After completing the telomere-to-telomere assemblies, we focused on characterizing the linear chromosomes, beginning with the telomeric ends, as these are the regions essential for chromosome stability and replication. We confirmed the completeness of the linear chromosomes via the presence of specific telomeric end motifs and by resolving the hairpin structures (Hepner et al. 2023; Tourand et al. 2009).

Our final objective was to investigate the genomes, as comparative genomics of closely related strains provides a powerful approach to understanding *Borrelia*’s virulence factors and genomic architecture in greater depth (Qiu et al. 2004). Since *B. burgdorferi* strains that cause Lyme disease vary in virulence, pathogenicity, and host-vector associations (Kurtenbach et al. 2002; Wolcott et al. 2021), and different *B. burgdorferi* isolates can have different infectivity (Wormser et al. 2008; Earnhart et al. 2005), the knowledge gained from this study will contribute to understanding these phenotypic variations in future studies. Additionally, the fully assembled genomes of Canadian endemic strains will serve as a major resource for the scientific community studying Lyme disease.

## Materials and methods

### *B. burgdorferi* culture and DNA extraction

Infected ticks were collected from four different sites in Canada: four sites in northwestern Ontario (Big Grassy, Big Island, Manitou Rapids, and Birch Island) and one site in Manitoba (Buffalo Point) (Supplementary Figure S1) during October and November 2016. The samples were obtained from the Public Health Agency of Canada (PHAC). Detailed procedures regarding tick collection, transport, storage of samples positive for *Borrelia burgdorferi*, and preparation of single-strain colonies are described in Tyler et al. (2018).

The nine *B. burgdorferi* strains were cultured in Barbour-Stoenner-Kelly (BSK-H) medium supplemented with 6% rabbit serum (Sigma, cat #B8291) in glass tubes at 33°C under 5% CO₂ without agitation for approximately 10 days. After this period, cultures were evaluated for spirochete count, morphology, and motility using dark-field microscopy (Laane 2013). DNA was extracted from the cultures using the Wizard® Genomic DNA Purification Kit (Promega, cat #A1120) following the manufacturer’s instructions. DNA was quantified using a Take 3 Cytation platform (Agilent), and its integrity and fragment sizes were assessed by low-concentration agarose gel electrophoresis.

### Library preparation and DNA sequencing

We performed whole genome sequencing using two different technologies, the Oxford Nanopore Technology (ONT) and Illumina (MiSeq) technology. Prior to library preparation, the DNA was quality controlled on a Genomic DNA ScreenTape (Agilent TapeStation). For short read sequencing, the libraries were prepared using the NxSeq AmpFREE kit (LCG) following manusfacturer’s instructions and controlled on a DNA ScreenTape (Agilent). The indexed libraries were multiplexed and sequenced Illumina MiSeq v2 flowcell in a paired-end 150 bp configuration. We obtained an average of 970,059 reads per sample with a 195x mean genomic coverage. For long read ONT sequencing, the gDNA size was assess using the FEMTO Pulse gDNA 165 kb Kit (Agilent) and the libraries were prepared following Oxford Nanopore instructions with native sample barcoding. The libraries were pooled and sequencing on two MinION flowcells (R10.3), obtaining an average of 78,515 reads per sample with an average 75x genomic coverage, median read length of 1,440 bp, N50 of 4,850 bp and median Phred score of 12.

### Assembly and annotation of the linear chromosomes

Quality control of the Illumina and ONT reads was performed using fastp (Chen et al. 2018) and Filtlong (https://github.com/rrwick/Filtlong) respectively. Nanoplot was used for visualization of the long read quality parameters before and after trimming (De Coster and Rademakers 2023). We used Trycycler (Wick et al. 2021) for assembling the linear chromosome using a hybrid long-read and short-read method as it allowed for more reliable consensus genome assembly (Supplementary Figure S2). We chose Canu, Flye and Raven as assemblers of choice in Trycycler. The assembled contigs were polished with both ONT and Illumina reads using Medaka (https://github.com/nanoporetech/medaka, Oxford Nanopore Technologies Ltd.) and Polypolish (Wick and Holt 2022), respectively. Due to the nature and length of plasmids in *Bb,* we observed significant fragmentation in plasmid assembly attempts with all selected assemblers. Therefore, we chose not to proceed with the consensus clustering step and opted for a more adapted approach described below in the plasmid assembly section.

We investigated the assembled contigs for terminal inverse duplication using ModDotPlot (Sweeten et al. 2024) and Inverted Repeats Finder (Warburton et al. 2004). After confirming the presence of terminal inverse duplications in both ends, which provides evidence that the contigs carry the full-length linear chromosome, including the hairpin loop structure, we looked for the exact inversion position in telomeric ends. Inversion points were identified based on the presence of the conserved right and left telomeric sequences of the reference strain B31 (Casjens 1999). We trimmed the contigs based on those inversion positions using a custom Python script. The length of the contigs before and after trimming are summarized in Table 1.

**Table 1.** Assembly and annotation metrics of the linear chromosomes of the nine strains.

| Strain | Length before trimming (bp) | Final length (bp) | CDS | Pseudogenes | Hypotheticals | GC (%) | tRNAs | rRNAs |
| --- | --- | --- | --- | --- | --- | --- | --- | --- |
| Bb16 122 | 931948 | 910513 | 812 | 0 | 13 | 28.6 | 33 | 5 |
| Bb16 131 | 932974 | 902779 | 811 | 2 | 13 | 28.6 | 33 | 5 |
| Bb16 133 | 918585 | 902780 | 811 | 1 | 13 | 28.6 | 33 | 5 |
| Bb16 134 | 924988 | 902780 | 811 | 1 | 13 | 28.6 | 33 | 5 |
| Bb16 22 | 925188 | 910197 | 816 | 0 | 15 | 28.6 | 32 | 5 |
| Bb16 33 | 933264 | 903010 | 810 | 0 | 13 | 28.6 | 32 | 5 |
| Bb16 49 | 930171 | 910024 | 813 | 0 | 14 | 28.6 | 32 | 5 |
| Bb16 47 | 926277 | 910705 | 814 | 0 | 15 | 28.6 | 32 | 5 |
| Bb16 57 | 919254 | 907757 | 812 | 0 | 13 | 28.6 | 32 | 5 |

The assembled and trimmed contigs were validated for accuracy by mapping the Illumina and ONT reads back to the final contigs using BWA (Jung and Han 2022) and Minimap2 (Liyanage et al. 2023), respectively (Supplementary Figure S3). Alignment outputs were filtered for secondary alignments using SAMtools (Li et al. 2009). The normalized coverage tracks were generated using DeepTools (Ramirez et al. 2014), and the coverage tracks were visually inspected using Integrated Genome Viewer IGV (Robinson et al. 2023). After validating the linear chromosome assemblies, gene annotation was performed using Bakta (Schwengers et al. 2021). The summary of the annotation is included in Table 1.

### Assembly and curation of plasmid

For assembling plasmids, we used Plassembler, a recently developed assembler specifically designed to assemble plasmids and tackles the challenges of assembling even the low copy number plasmids (Bouras et al. 2023). It assembles the linear chromosome separately using a hybrid method and then maps the ONT and Illumina reads back to the assembled linear chromosome to remove the chromosomal reads and perform the assembly using the remaining reads, assuming the remaining reads are mostly coming from plasmids. Plassembler generated hundreds of contigs for each strain. However, we selected the contigs that are at least 1000 bp as potential plasmid contigs. Plassembler compared each contig against the PLSDB plasmid database (Galata et al. 2019) as part of its pipeline to identify the closest plasmid orthologs in the database for annotation purpose and also perform the initial circularity test.

During the post-processing of the Plassembler’s output, we kept one contig per plasmid type for each strain. If there are two contigs, we selected the complete assembly over the partial assembly. If completeness is not confirmed for both contigs by Plassembler, we checked the identity of the partial assemblies with the reference using BLAST and then picked the one with longer alignment with the reference among the two partial assemblies. Contigs that are less than 20% in size compared to their corresponding reference were discarded from the downstream analysis. Contigs that are at least 50% of the reference plasmid in length but we could not confirm completeness using the downstream plasmid completion pipeline (Supplementary Figure S5) were classified as partial.

### Completion of circular and linear plasmids

Plassembler uses Unicycler’s (Wick et al. 2017) graph-based method for detecting overlapping ends of each contig and trimming it on one end to generate a complete assembly of the circular plasmids. However, Unicycler might fail to identify an overlap and circularize a contig, especially for complex *Borrelia* plasmids which carry high sequence similarity in distant region of the contig (Casjens et al. 2017). In that case, we investigate the contigs for direct overlapping ends using self-identity dot plots created by ModDotPlot (Sweeten et al. 2024). If terminal direct overlaps are identified, we removed the overlapping sequences at one end using Simple-Circularize (Kzra 2018). Finally, we aligned ONT reads back to the trimmed circular plasmids using minimap2 (Li 2018). SAMtools was used to convert alignment output and create an index file (Li et al. 2009). The coverage of ONT reads for each complete circular plasmid was investigated to verify the accuracy of the assembled contigs. Terminal unmapped reads at both ends of the circular plasmids visualized by IGV confirmed the circularity of the final trimmed circularized contigs.

For linear plasmids, most contigs generated by Plassembler did not contain terminal inverse duplications. However, to confirm the completeness of linear plasmids, the contigs need to carry the telomeric hairpin wraparound indicated by the terminal inverse duplication (Hepner et al. 2023). To solve this issue, we extended the Plassembler assembly pipeline (Supplementary Figure S5A) and customized it for assembling *Borrelia* linear plasmids. In summary, we extracted the ONT reads that map to the terminal 5000 bp of each linear plasmid contig using SAMtools (Danecek et al. 2021), and BEDtools (Quinlan 2014). After that, we assembled those extracted reads from both ends using a long-read assembler Flye (Freire et al. 2022), which generated two telomeric end contigs, one for the left end and the other for the right end. We then merged these two telomeric end contigs with the plasmid contig using BLAST (Altschul et al. 1990) and an in-house Python script. In the Python script, we recorded the overlapping alignment positions from each pairwise BLAST result, trimmed the overlapping sequences from the terminal end contigs, and only concatenated the unique sequences of each terminal end contig to the plasmid contig. In the end, the concatenated plasmid contigs were each investigated using ModDotPlot (Sweeten et al. 2024) for terminal inverse duplication at both ends (Supplementary Figure S5B). If inverse duplications were present, we trimmed the concatenated plasmid contigs based on the inversion positions identified by Inverted Repeats Finder (Warburton et al. 2004). If a single or no inverse duplication were detected, we classify those contigs as partial assemblies. The final assembled complete and partial plasmids were annotated using Bakta (Schwengers et al. 2021) and the annotations were visualized using Prokase (Grant et al. 2023).

### Phylogenetic analysis of the core genome

Publicly available *Borrelia burgdorferi* genome assemblies (n=445) were downloaded from NCBI (Sayers et al. 2024) to perform a phylogenetic analysis, including our nine newly assembled full genomes. Among the 445 genome assemblies, we have geographical locations available for 418 assemblies. First, we downloaded the metadata of the 445 assemblies using the Bioproject IDs from NCBI. We extracted the Biosample IDs and the geographical locations of assemblies from the metadata (Supplementary Data 1). Among the 445 assemblies, 121 assemblies had an assembled contig for linear chromosome that is at least 0.8 Mb in length. We extracted those 121 linear chromosomes and with our assembled linear chromosomes, a total of 130 linear chromosomes were used for phylogenetic analysis. All assemblies were converted to forward strands before running alignment using MAFFT (Katoh et al. 2019). A maximum likelihood-based phylogenetic tree was generated using the iqTree approach with 1000 bootstraps (Minh et al. 2020). The phylogenetic tree was visualized using iTOL (Letunic and Bork 2024).

### Comparative genomics

The nine assembled genomes were aligned to the reference genome B31, and a consensus sequence was created using MAFFT (Katoh et al. 2019), masking the variable regions across the linear genome. The nine assembled and the reference B31 aligned linear chromosomes were compared to the consensus base-by-base, and every mismatch position compared to the consensus was recorded. A mismatch visualization plot was created using the CMplot (Yin et al. 2021) package in R. The variable regions of the nine assembled genomes were investigated using MAUVE (Darling et al. 2004). The right end of the telomeres (from 900 kb to the end) of each assembled genome was investigated using BLASTn, and a circular alignment plot was created using Circuletto (Darzentas 2010).

A plasmid similarity matrix was computed to assess the overall plasmid similarity between the assembled strains and reference strains (297, B17/2013, B31, B331, B31_NRZ, B500, JD1, MM1, MN13-1420, N40, ZS7). For each strain, we quantified the fraction of assembled plasmids orthologous to a specific reference strain using the following formula:

Similarity score of strain X against reference Y = Number of assembled plasmids orthologous to plasmids in strain Y / Total number of assembled plasmids in strain X

After computing similarity scores for all strains, we generated a clustermap to visualize strain relationships based on their plasmid profiles.

## Results

### Assembly of telomere-to-telomere *B. burgdorferi* linear chromosomes

Telomere-to-telomere complete genome assemblies were generated for the nine *Bb* strains isolated from infected ticks collected across multiple sites in Canada. These included four sites in northwestern Ontario (Big Grassy, Big Island, Manitou Rapids, and Birch Island) and one site in Manitoba (Buffalo Point) (Supplementary Figure S1). In addition, we developed a genome assembly pipeline tailored to complex linear chromosomes and plasmids of *Bb* genomes which can be broadly applicable to other *Borrelia* species.

One of the main challenges to assembling *Borrelia* linear chromosome and plasmids is to assemble it up to the telomeric end due to the presence of a hairpin wraparound at each end. To overcome this challenge, we used ONT long-read sequencing, which could capture the terminal inverse duplication in a single read (Figure 1A). For the linear chromosomes, the minimum, maximum and median ONT coverage at the hairpin telomeres are 12x, 300x, and 29x respectively. Therefore, Trycycler could assemble the telomeric inverse duplication for all strains without requiring an individual assembly approach for the telomeres. Using a self-identity dot plot approach (Figure 1B), the inverse duplication at both telomeres were detected, ranging from 5 to 18 kb in length, with the exception of the right telomeric end of isolate Bb16-133, where only a shorter inverse duplication (68 bp) got assembled. Length of the assembled linear chromosome contigs before and after trimming the inverse duplication at the telomeres, as well as number of annotated genes, are reported in Table 1.

**Figure 1.**
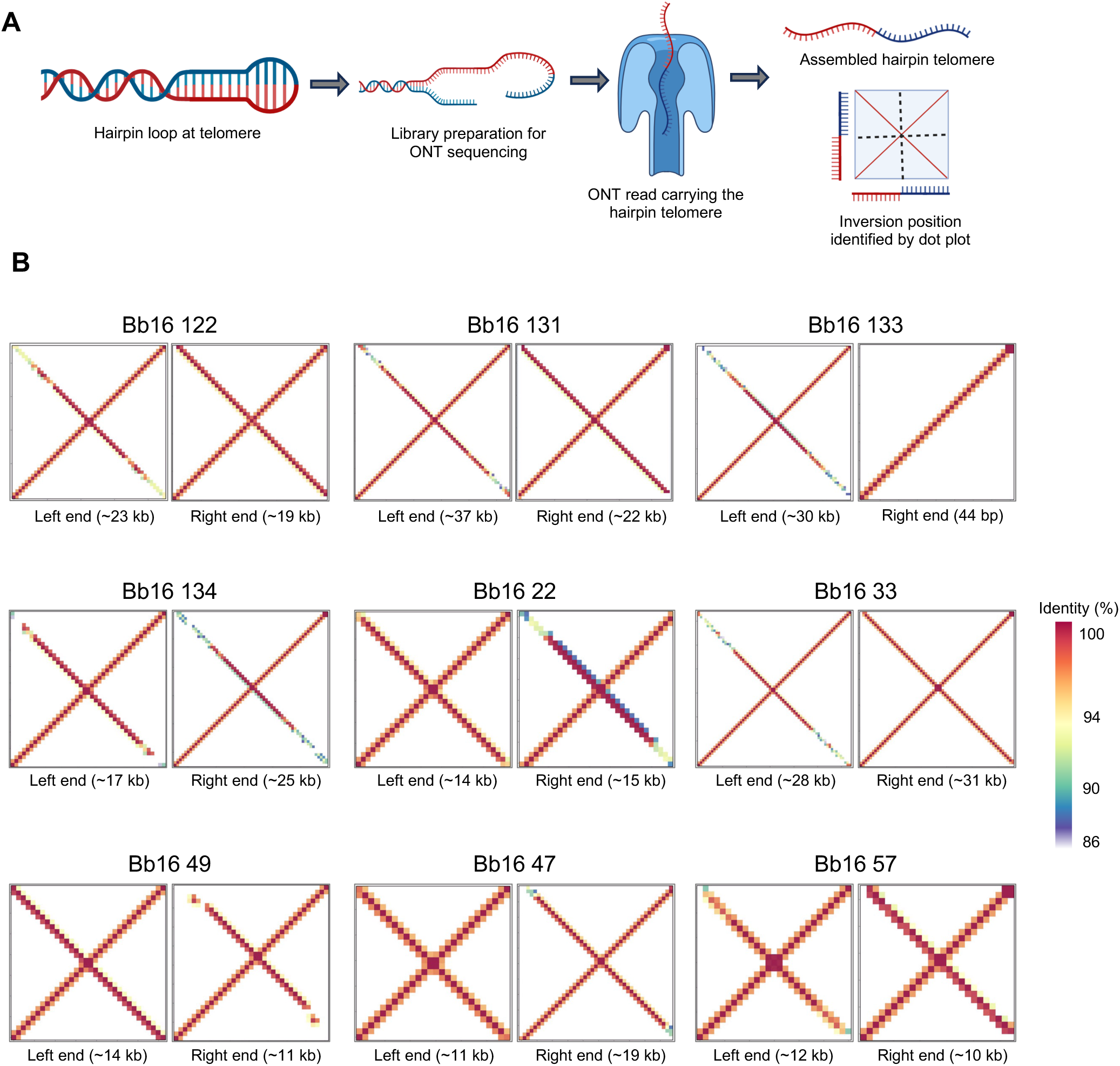
Hairpin telomeres identified in the linear chromosome of the nine strains. **A)** Schematic representation of the identification of *Borrelia* hairpin telomeres using Oxford Nanopore Technology long-read sequencing. **B)** Dot plots showing the telomeric hairpin structures as inverted duplications at both ends of the linear chromosomes for the nine assembled strains. The length of the assembled inverted duplications is indicated in brackets.

It is important to mention that we did not have Illumina coverage at the wraparound sequences since the folded hairpins prevent adapter ligation and PCR amplification. Therefore, ONT reads were essential for telomeric wraparounds assembly and validation. Each side of the hairpin loop sequences were consistent, as in most cases both forward and inverse sequences carry the same genetic variation compared to the reference B31 (Figure 2), indicating the applicability and accuracy of the ONT sequencing for assembling the telomeric hairpin loop structure.

**Figure 2.**
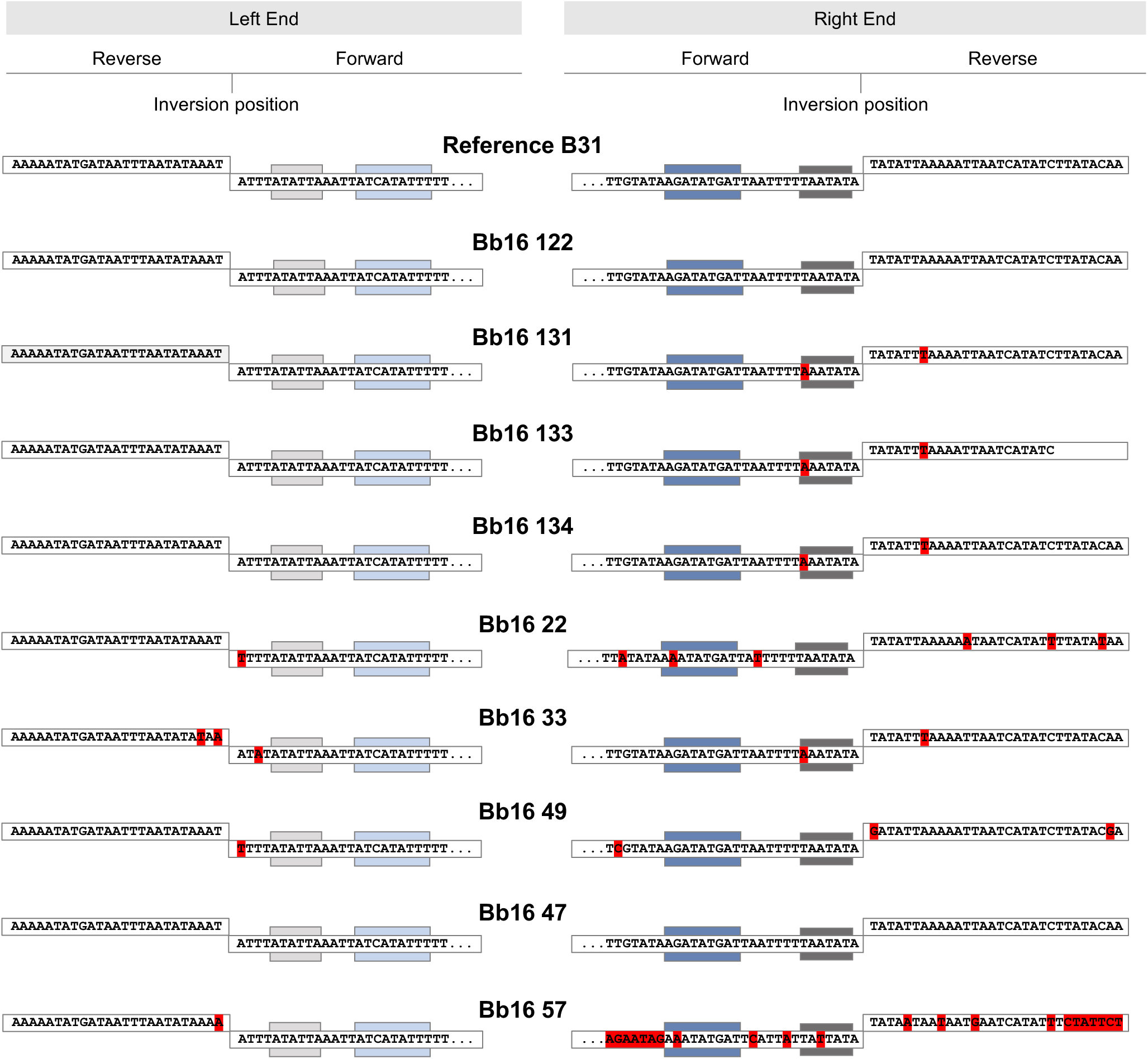
Conserved telomeric motifs detected at inversion points of both chromosomal ends in all strains. Forward and reverse alignments with the reference B31 telomeric sequences are shown as clear boxes. Polymorphisms relative to the reference are highlighted in red for single nucleotide polymorphisms (SNPs). Conserved telomeric motifs identified in the telomeric sequences are indicated by grey and blue boxes, where darker shades represent the reverse complements of the identified motifs.

### Genome assemblies revealed a diversity in conserved telomeric sequence

Although we successfully assembled the telomeric wraparound for all strains, identifying a precise approach to trim the linear chromosomes, ensuring that the wraparound sequences appear only once in the final linear chromosomes, proved challenging. Since the conserved telomeric end sequences of the reference strain B31 are characterized, except for the extreme one or two nucleotides (Casjens 1999), we searched for the 30-32 bp reference conserved sequences at the wraparound region of the assembled contigs. Our goal was to trim the linear chromosomes precisely at the inversion point where the conserved telomeric sequences align in both the forward and reverse orientations, while also comparing the conserved sequences of the assembled strains to the reference B31.

Both the right and left conserved telomeric sequences of the reference strain B31 were found in the chromosome assemblies of all strains, allowing us to pinpoint specific inversion positions (Figure 2). We confirmed the nucleotides of the extremity of hairpin wraparound in all strains. We observed that the left telomeric sequence is more conserved than the right. Across the nine strains, Bb16-22, Bb16-33, Bb16-49, and Bb16-57 exhibited SNPs on the left telomeric end, while the remaining strains are 100% identical to the reference. In contrast, the right telomeric sequences showed greater variability, where among the nine strains, only Bb16-122, Bb16-47 carries the reference conserved telomeric sequence with 100% identity, while the others carry substitutions; Bb16-57 is only 56% identical to the reference. Interestingly, most of these variants are located outside of the conserved motif ATATTA, ATCATYTNT (Huang 2004) (Figure 2). Among these motifs, ATATTA is located one to four nucleotides from the terminus, and ATCATYTNT is located 14 bp from the end and is found in all previously known *Bb* telomeres (Huang 2004). The presence of these two motifs in all assembled strains provides evidence that all nine strains harbor a Type 1 telomere (Huang 2004). Despite all strains carrying the same telomere type, we identified two variants of the reference conserved motif TATAAT: TATAAA and TATTAT at the right telomeric sequence. Among the nine strains, Bb16-131, Bb16-133, Bb16-134 and Bb16-33 carry the same variant TATAAA, Bb16-57 carry the variant TATTAT, further indicating the diversity within the conserved telomeric sequences of these *B. burgdorferi* strains (Figure 2).

### Phylogenetic analyses show genetic diversity within and outside Canada

To evaluate the genetic similarity among the assembled strains, we performed an SNP-based phylogenetic analysis, comparing the conserved sequence of the linear chromosomes of the assembled strains to the reference B31. The result shows that the three strains from Manitoba exhibit high similarity in their SNP profile compared to the reference, with Bb16-49 and Bb16-57 being the most closely related and clustering together (Figure 3A). However, there is more diversity among the strains from Ontario, as we saw the six strains are not monophyletic. The three strains from Big Island and the one from Birch Island formed one cluster, while those from Manitou Rapids and Big Grassy were found to be most closely related to the reference strain B31 (Figure 3A).

**Figure 3.**
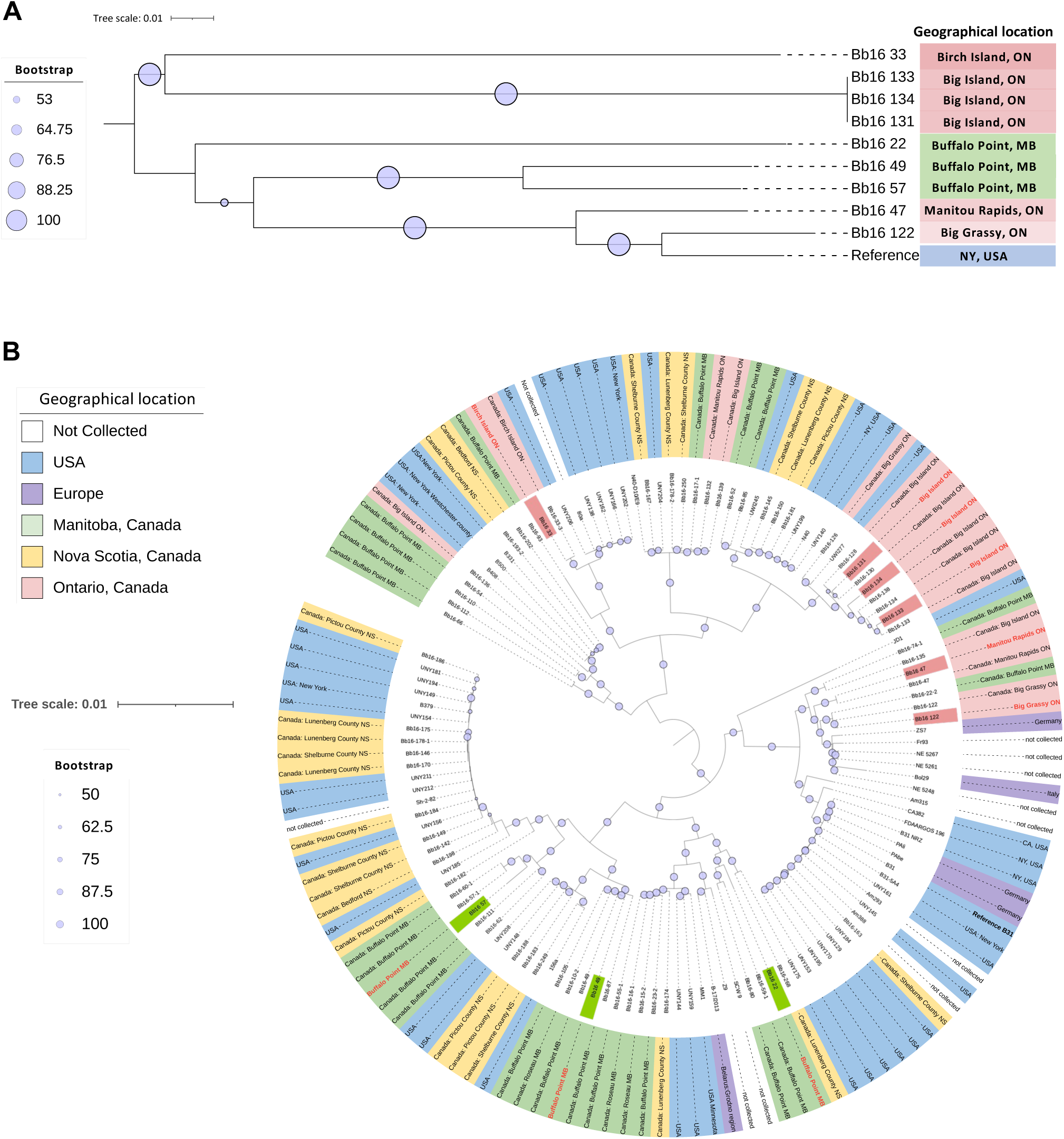
Phylogenetic analysis of the nine newly assembled Canadian strains based on the linear chromosomes. **A)** SNP-based phylogenetic tree illustrating the relationships among the nine assembled genomes; their geographical locations and bootstrap values are indicated. SNPs were identified by comparison to the reference strain B31. **B)** Maximum likelihood phylogenetic tree including 121 publicly available *B. burgdorferi* genomes from various geographic regions along with the nine newly assembled Canadian strains. The tree was constructed using IQ-TREE v2.3.5 with 1,000 bootstrap replicates to assess branch support.

To investigate the genetic diversity among the *Borrelia* strains within and outside Canada, we created a maximum likelihood phylogenetic tree. This analysis included a total of 70 Canadian strains coming from three provinces, Ontario, Manitoba and Nova Scotia, including the nine newly assembled strains, along with additional strains from the USA and Europe. The results indicate that the linear chromosomes of *Borrelia* across different geographical locations are well conserved and co-linear with the reference, which was reflected by the small branch length (<0.005 expected substitutions per site) in the phylogenetic tree. However, the slight genetic differences we detected do not correlate well with the geographical location (Figure 3B). We did not see any distinct clustering within the Canadian strains among the three provinces. The three assembled strains from Manitoba and six strains from Ontario belong to separate lineages, each containing samples from the USA or from other distant Canadian provinces. The overall diversity among the linear chromosomes of the Canadian strains also suggests the rapid migration of *Borrelia* across the provinces and the USA, which might be facilitated by the migration of the vector or the animal host across the provinces.

### Variability of linear chromosomes originates from the right telomeric end

The linear chromosomes of the nine strains are largely syntenic with high sequence identify and little to no variation in nucleotide composition (e.g., SNPs or structural rearrangement) for ∼92-93% of their total length. This region of the linear chromosome is defined as core conserved region, totalling ∼847 kb in size in the assembled strains (Figure 4A). Comparison of the nine assembled strains and the reference B31 with the core consensus sequence, which was generated from the core conserved regions of the nine linear chromosomes and the reference, shows an accumulation of SNPs at the right end of B31 linear chromosome, highlighting a notable difference between these Canadian strains and the current reference (Figure 4A). This finding suggests that future studies on Lyme borreliosis in Canada, particularly those studies focusing on SNP analysis, should consider using the newly assembled strains as a reference instead of B31.In addition, the differences within the Canadian strains in terms of the SNPs in this region reflects the phylogenetic diversity observed before (Figure 3A) and highlights the diversity of these assembled strains based on their core conserved linear chromosomes at the right end (Figure 4A).

**Figure 4.**
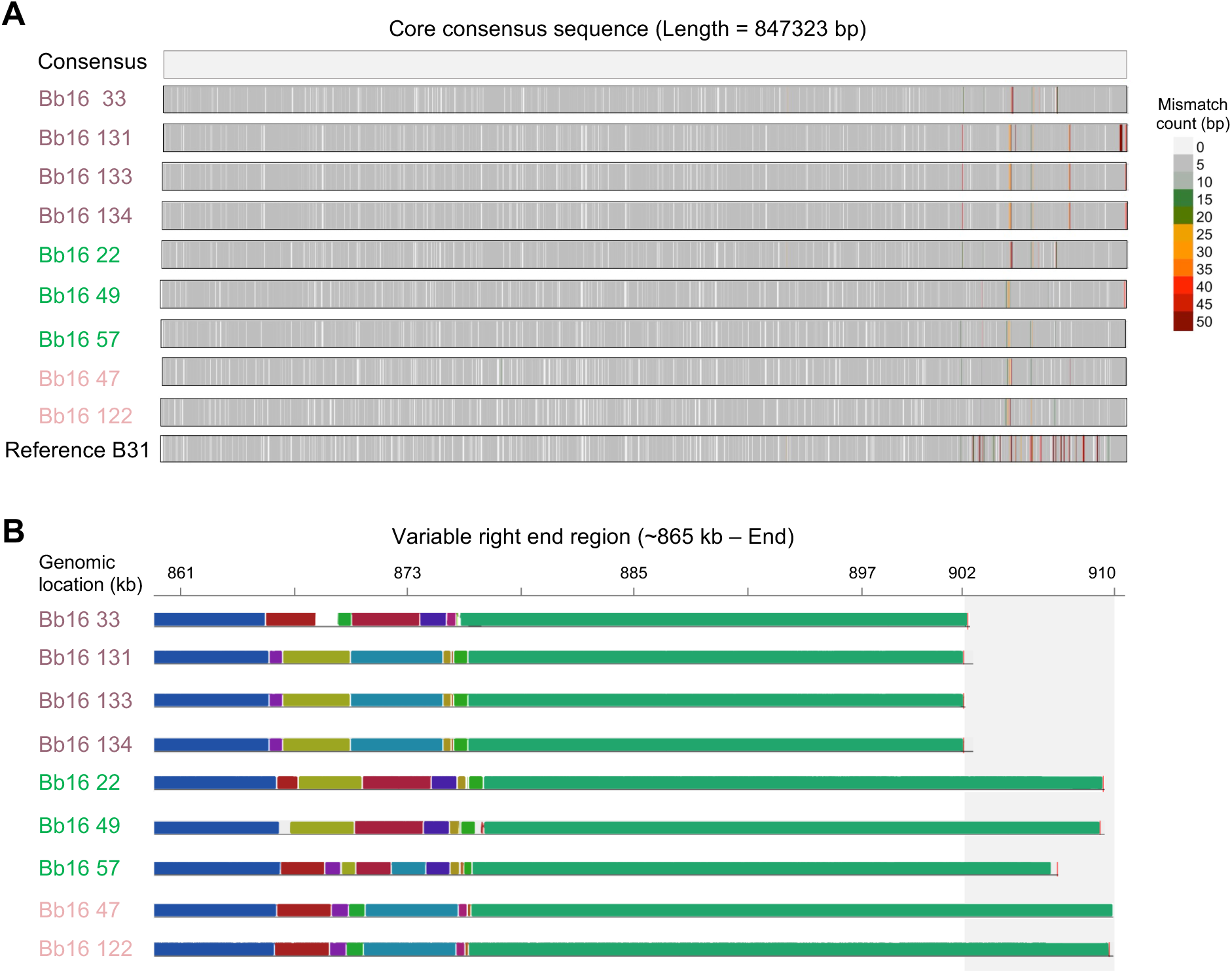
Genetic diversity of the nine strains at the conserved and variable regions of the chromosomes. **A)** Single nucleotide mismatches across the conserved regions of the linear chromosome, calculated in 50 bp windows, by comparison with a consensus sequence generated from the nine assembled genomes and the reference strain B31. **B)** Visualization of genetic rearrangements and variability in right telomeric end lengths using MAUVE alignment. The names of the assembled strains are colored according to their geographical locations (green = Manitoba; and pink shades = Ontario).

In addition to examining the core conserved sequence similarity, we investigated the variable regions, which begins at ∼865 kb in the assembled strains (Figure 4B). We observed that in four strains (Bb16-33, 131, 133, and 134), the linear chromosome extends to approximately ∼902 kb, whereas in the remaining strains, it is 5 to 10 kb longer, indicating a high level of variability at the right telomeric end. While previous studies reported variability within the last ≤ 20 kb of the right chromosomal end (Casjens et al. 2012), we identified a region with genetic rearrangements from 865 to 877 kb in the assembled strains (Figure 4B). Interestingly, the pattern of genetic rearrangement in this genomic segment (from 865 to 877 kb) matches the SNP-based observation in the core conserved sequences (Figure 4A, B). Notably, Bb16-33 and Bb16-57 exhibit the most distinct genetic rearrangement among the nine strains. We identified three other regions of genetic variability across the linear chromosomes at ∼213-215 kb, ∼557-559 kb, and ∼845-847 kb (Supplementary Figure S4). These findings reinforce the notion that the assembled Canadian strains differ significantly from the current reference strain B31 over multiple regions. It also supports the fact that closely related *Borrelia* strains evolve not only through point mutations, but also via genetic rearrangements at the variable regions (Schutzer et al. 2011).

### Telomeric extension at the right end carries a genomic segment of plasmid lp28-1

The linear chromosome of the *Borrelia* genome has been previously known to be highly variable at the right end, where this variation can result from the insertion of plasmid-like DNA (Casjens et al. 2012). In the reference strain B31, this variable region starts at 903 kb and is >98% identical to the linear plasmid lp28-1 of strain 297 (Casjens et al. 2012). In our assembled strains, we observed that five strains − Bb16-122, 47, 22, 49, and 57 − contain a ∼5-8 kb right end extension beyond the right-end position of the other four strains, which terminate around ∼902 kb (Figure 4B). This observation prompted us to investigate the source of these extensions. We found that the right end extensions of our assemblies have high similarity with a linear plasmid lp28-1 of the strain 64b (Accession no. CP001423.1). All strains carried a segment of different length from DNA sequence >98% identical to the first ∼10 kb of the plasmid lp28-1 of 64b strain, extending from ∼900 kb to the chromosome’s end (Figure 5). The length of this homologous segment varies among strains: Bb16-131, Bb16-133, Bb16-134, and Bb16-33 contain a shorter segment (∼2.4 kb), whereas Bb16-122 and Bb16-47 showed the largest alignment of ∼9.5 kb, which carries 12 complete genes from plasmid lp28-1 (Figure 5). Among these genes, four are annotated as *argF* (ornithine carbamoyltransferase), *BBU64B_F0011* (arginine-ornithine antiporter), *BBU64B_F0009* (putative lipoprotein), and *BBU64B_F0002* (type I restriction endonuclease), while the remaining are conserved hypothetical protein. This result indicates that the genomic plasticity of *Borrelia* at the right end might contributes to the difference in gene content among even the closely related strains, as previously suggested by (Casjens et al. 2012). The fact that strain 64b was collected from NY, USA (Schutzer et al. 2011), also raises the possibility of a recent common ancestor contributing to the hybrid telomeric end sequences in the Canadian strains.

**Figure 5.**
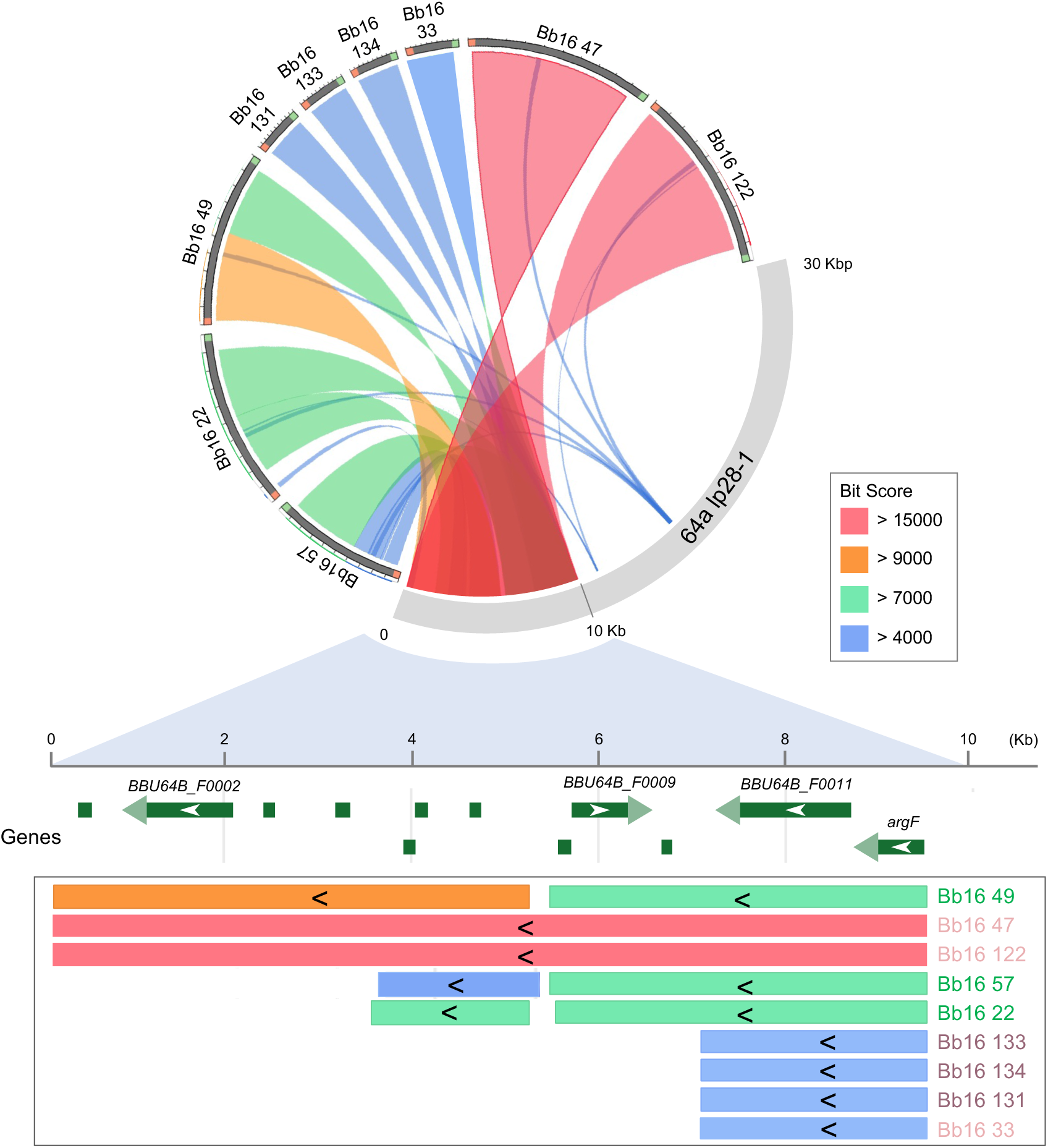
Telomeric end length variation among the nine assembled strains is driven by the insertion of a plasmid-like DNA segment. A terminal segment of variable length (∼2-10 kb) from the linear plasmid *lp28-1* of strain 64b was identified at the right telomeric ends of all nine linear chromosomes. The alignments showed 99% sequence identity, in reverse orientation, and are colored based on the alignment bit score. Genes located within the shared *lp28-1* segment and their corresponding positions in the chromosome ends of each strain are indicated. Strain names are colored by geographic origin (green = Manitoba; pink shades = Ontario).

### Complete assembly of linear and circular plasmids

One of the challenging aspects of *Borrelia* genome assembly is to assemble the complete set of plasmids, especially for the linear plasmids. Although draft genome assemblies were available for 64 Canadian strains, the complete plasmid assemblies were lacking. Therefore, a primary goal of this study was to develop an optimized pipeline to assemble both circular and linear plasmids. The pipeline we developed (described in the methods section) (Supplementary Figure S5) utilizes two assemblers, Plassembler and Flye (Bouras et al. 2023; Freire et al. 2022), and effectively assembles plasmids in a relatively straightforward approach.

For circular plasmids, we achieved complete assembly using Plassembler, followed by additional steps of contig circularization and trimming validation (Supplementary Figure S5). An example of this trimming process is shown for cp32-12 contig (Supplementary Figure S5B), where validation of terminal direct duplication removal was confirmed by a self-identity dot plot. Assembly of the linear plasmids on the other hand was more complex since Plassembler failed to completely assemble most of the linear plasmids. To address this limitation, we extended our pipeline with a “Telomeric End Extension” method (Supplementary Figure S5A), which enabled complete assembly of most linear plasmids. We successfully identified and trimmed the terminal inverse duplication for most contigs. An example of the trimming process is provided for the linear plasmid lp38 (Supplementary Figure S5B). However, for a few contigs, the telomeric inversions in both ends were not detected, thus we classified them as partial. Examples of these partial plasmids include lp28-5 of Bb16-49, lp28-3 of Bb16-22, and lp28-2 of Bb16-13, where the left telomeric end inversions were absent.

In total, we assembled between 10 and 15 plasmids for each strain (details in Table 2). Overall, this collection includes 14 circular plasmid types and 15 linear plasmid types. Notably, two conserved plasmid types, cp26 and lp54, were assembled for all nine strains (Table 2). Since the cp26 and lp54 plasmids were assembled in all strains, we further investigated the *ospA*, *ospB* (located on plasmid lp54), and *ospC* (located on cp26) gene types. By comparing the assembled *ospC* gene sequences to the reference *ospC* types as defined by a previous study (Russell et al. 2024), we classified our strains into 7 types (A, B, C, D, F, H, and N), all of which are infectious *ospC* types (Russell et al. 2024) (Figure 6A). Interestingly, strains Bb16-131, 133, and 134 belong to the same *ospC* type, providing additional evidence of their close genetic relationship. It is worth noting that *ospA* is a key antigen targeted by Lyme disease vaccine (e.g., LYMErix), and continues to be a focus for developing new vaccines (Dattwyler and Gomes-Solecki 2022). Phylogenetic analyses of the *ospA* and *ospB* genes clustered the strains into two distinct groups, highlighting the diversity of these genes even among strains from the same geographical locations (Figure 6A).

**Table 2.**
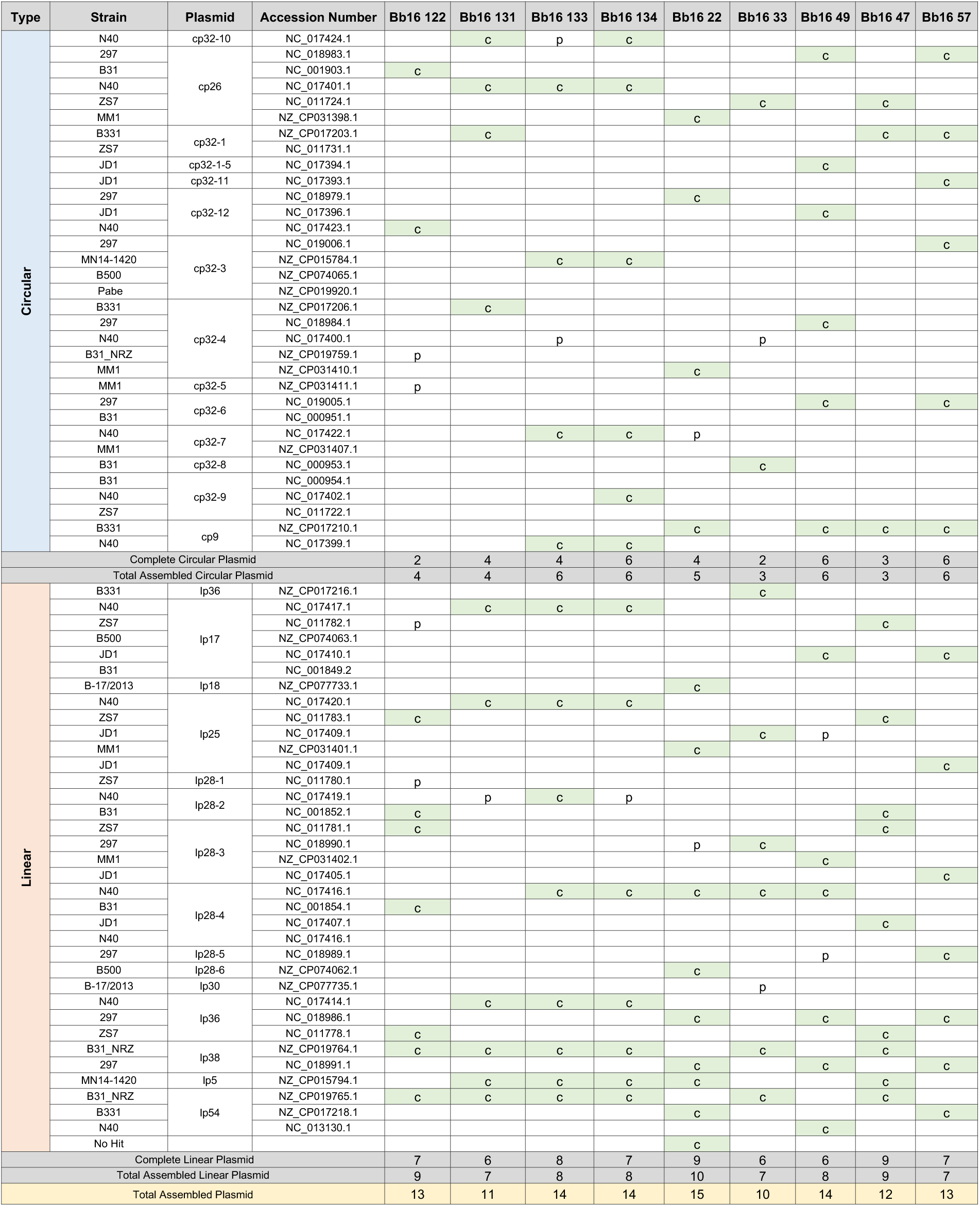
Summary of the assembled linear and circular plasmids in the nine Canadian strains.

**Figure 6.**
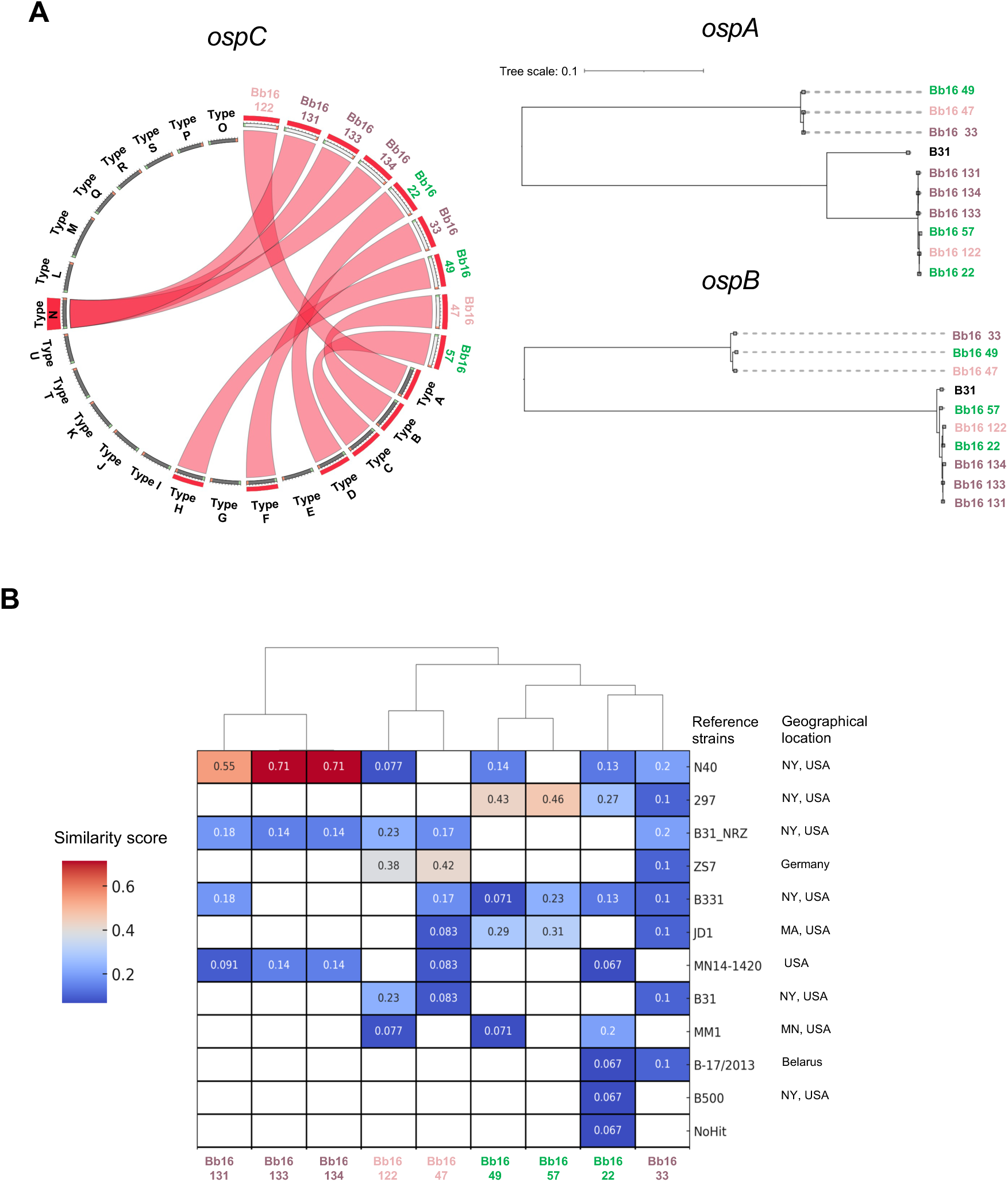
Genomic diversity among the nine assembled strains is reflected in their plasmid profiles and key surface protein genes. **A)** Diversity is observed in the *ospC* type of the nine strains. Phylogenetic trees based on the *ospA* and *ospB* genes also show diversity among the strains. The names of the strains are colored according to their geographical locations (green = Manitoba; pink shades = Ontario). **B)** Plasmid similarity scores were calculated for all strains based on the percentage of assembled plasmids matching to specific reference strains.

In addition to assembling the conserved plasmids, linear plasmids lp36 and lp38 were assembled in eight and nine strains respectively, making them the two most frequent linear plasmids in the assembled strains. Notably, strain Bb16-22 exhibited the highest number of assembled plasmids, 5 circular and 10 linear, including a new plasmid subtype (Supplementary figure S6). Therefore, we recommend this strain as reference for future studies on Canadian *Borrelia*. Its extensive plasmid repertoire provides a valuable resource for comparative genomics and evolutionary studies.

### Diversity of plasmid profiles in the assembled strains

A clustermap based on plasmid similarity scores across all assembled strains revealed two distinct groups, similar to the pattern observed above for linear chromosomes (Figure 3A). The three strains from Ontario Bb16 131, 133, and 134 and two strains from Manitoba Bb16 49 and 57 display similar plasmid profiles. (Figure 6B). Among 11 reference strains containing plasmids, N40 has the most similar profile to strains Bb16-131, Bb16-133, Bb16-134, while strain 297 is most similar to the two Manitoba strains Bb16-49 and Bb16-57. Interestingly, both N40 and 297 strains were isolate in New York, USA. The plasmid profiles of Bb16-33 and Bb16-22 are the most divergent, as indicated by their evenly distributed similarity scores across reference strains. Another key observation is that assembled and reference strains identified as similar in core genome-based phylogenetic analyses (Figure 3B) also share similar plasmid profiles. This suggests that plasmid diversity is generally correlated with core genome diversity.

### Identification of a novel lp28-type linear plasmid in strain Bb16-22

Among the 10 linear plasmids assembled in strain Bb16-22, one did not match any hit in the PLSDB (Galata et al. 2019), preventing Plassembler from assigning a reference ortholog. To rule out the possibility of a misassembly, we conducted a thorough evaluation of the following criteria: 1) the contig had full and uniform coverage of ONT reads, 2) the contig had unmapped overhanging ONT reads at both ends, 3) the contig had the terminal direct or inverse duplications in both ends before trimming, 4) the contig has conserved telomeric sequences and motifs at both ends, and 5) the contig carry the plasmid maintenance, replication and partitioning genes which are required for the plasmid stability. This new Bb16-22 contig fulfilled all these criteria, confirming that it represents a valid linear plasmid (Figure 7A-C). In addition, a Pfam database search (Finn et al. 2014) identified protein families characteristic of *Borrelia* plasmids, including *Oms28* porin, *Borrelia* lipoprotein, and variable surface antigen *vlsE*. Since *Borrelia* plasmids are named according to their size and characteristic genes, and this plasmid is ∼28 kb in size with a characteristic *vlsE* gene cluster, we classified it as a novel lp28-1a subtype plasmid. However, a BLASTn search revealed that this new plasmid does not have a single best hit with other lp28 type plasmids. Instead, we found identical DNA sequences scattered across six other known linear plasmids (lp25, lp28-3, lp38, lp28-2, lp36, and lp28-1), with coverage ranging from 5 to 30% and sequence identity up to 90% (Figure 7D). Notably, lp28-1 contributes to at least 35% of the new lp28-1a plasmid’s sequence.

**Figure 7.**
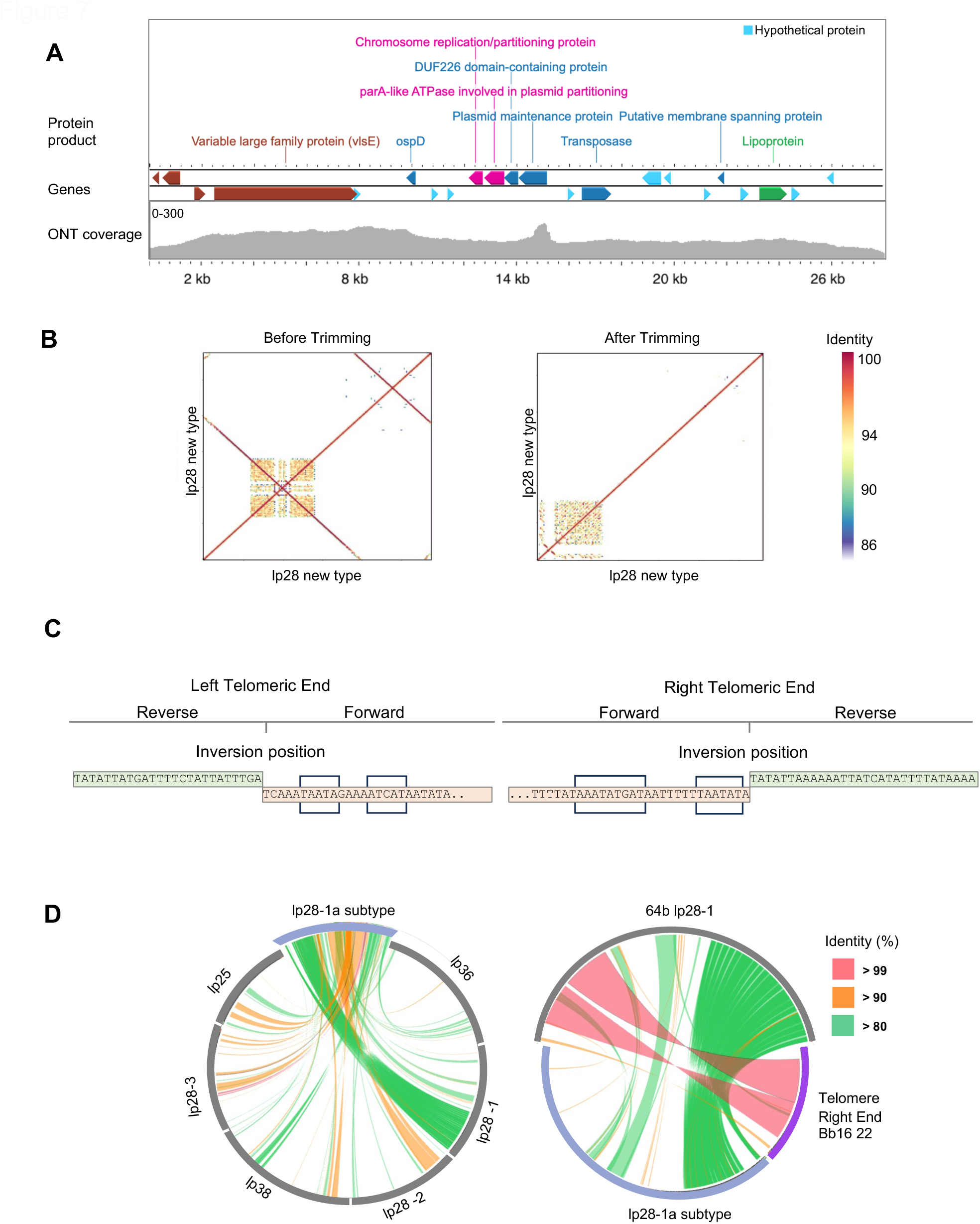
Identification of a novel fusion plasmid, lp28-1a subtype, in strain Bb16 22. **A)** ONT read coverage across the plasmid lp28-1a subtype. Annotated genes are depicted, including the *vlsE* antigenic variation cassette region. **B)** Detection and trimming of hairpin telomere wraparounds at both ends of the plasmid. **C)** Conserved telomeric sequences along with the conserved motifs, are present at both telomeric ends of lp28-1a. **D)** The novel plasmid was formed through the fusion of genomic segments from six different linear plasmids and the remaining DNA from the *lp28-1* plasmid, resulting in the unique *lp28-1a* subtype.

We showed that the linear chromosome of our Canadian strains carries a 2-10 kb DNA segment from the plasmid lp28-1 at their right telomeric end (Figure 5). Interestingly, we were unable to assemble a complete lp28-1 plasmid for any of the strains (Table 2), raising the question of whether the transfer of lp28-1 DNA to the chromosome telomere affects the plasmid stability and what segment of lp28-1 is integrated into the new lp28-1a plasmid. When we analyzed the composition of the DNA segment in the new lp28 plasmid and its similarity with the strain 64b lp28-1 and the right-end sequence of Bb16-22 linear chromosome using BLASTn, we observed that one end of the lp28-1 contributed approximately 10 kb into the right end of the linear chromosome, while the DNA segments from the opposite end got integrated into this new plasmid (Figure 7D).

## Discussion

### The developed assembly pipeline reduces manual intervention

Assembling complete *Borrelia burgdorferi* (*Bb)* genome is a challenging task that requires an efficient assembly pipeline particularly tailored toward fragmented genome combining linear and circular components. An ensemble pipeline, highlighting the difficulty of assembling the complete *Bb* genome using a single assembler was published where the approach relied on three different assemblers and used a consensus strategy to create the final assembly (Hepner et al. 2023). This workflow requires extensive manual inspection of numerous contigs generated by the three assemblers at several quality control and refinement steps, as well as during generation of the final consensus. To reduce the level of manual intervention throughout the assembly process, we developed a new pipeline that also leverages multiple assemblers but in a more streamlined manner (Supplementary Figure S2 and S5). By separating the assembly of the core chromosome from plasmids, we limited the refinement steps to the plasmids only, resulting in a more efficient process in terms of time, manual intervention, and accuracy. We adopted the Trycycler approach (Wick et al. 2021), enabling the use of three different assemblers as suggested by Herper et al. 2023 but with minimal manual investigation of contigs and concatenation steps. Instead, Trycycler itself handled the consensus generation steps, producing a final consensus linear chromosome. However, Trycycler is not designed for and did not produce high quality plasmid contigs. Therefore, for plasmid assembly we chose a separate assembler, Plassembler, specifically designed to capture even the low copy number plasmids (Bouras et al. 2023). This approach reduced the complexity of working with multiple contigs per plasmid, which is labor-intensive and prone to errors due to the high degree of sequence similarity among some *Bb* plasmid types. While Plassembler initially struggled to assemble linear plasmids, as trimming linear plasmids is not part of its pipeline, extending the pipeline with a contig trimming and refinement method significantly improved its performance. Overall, the pipeline we developed successfully produced complete assemblies for nine strains with minimal manual intervention and high efficiency. This pipeline is broadly applicable to other complex *Borrelia* genome, especially for the linear replicons. The detail pipeline including the codes are available as a STAR Protocol article.

### Comparative analysis of telomeres reveals major variations

Advancement in sequencing technology and the development of sophisticated assemblers resulted in improvement of *Bb* genome assemblies. From 2021 to 2025, 27 complete reference *Bb* genome assemblies were submitted to NCBI, including five strains from Canada. Most of these genomes were assembled using long-read sequences. However, it is unclear whether the telomeres of the linear replicons in these genomes are complete since detailed published methods and analyses are missing for most of them. While the increase in available assemblies is promising, it is important to confirm their completeness before investigating different aspects of the *Bb* genomes to uncover their complexities and their relationship with the pathophysiology of Lyme disease.

One of the most intriguing features of *Bb* genomes is their hairpin telomeres. Despite the fact that the genome assemblies of several *Bb* strains are reported as complete, the sequence of the telomeres for most of them are unknown or improperly validated due to the difficulties in sequencing the covalently closed hairpin ends (Tourand et al. 2009). Therefore, as our primary goal was to develop a pipeline to fully assemble, telomere-to-telomere, Bb genomes, we emphasize our approach to assemble and validate the telomeres of the linear replicons with high confidence and then investigate them more closely. In addition to reporting the 30 bp conserved sequence at the right telomeric end and the 25 bp conserved sequence at the left telomeric end of the nine Canadian *Bb* genomes, we detected variations in these conversed sequences, particularly on the right end. Notably, the single base variations on the left end occurred at the hairpin’s center where the inversion begins, and similar mismatches were observed for two strains at the right end (Figure 2). This type of terminal nucleotide mismatches in *Borrelia* telomeres is unusual since most *Bb* telomeric ends where the hairpin sequences are known are perfect palindromic (Casjens 1999). These mismatches could be explained by steric constraints that prevent the terminal 2-3 nucleotides from forming a double helix (Casjens 1999). Nevertheless, the sequence variations identified in this study raise the question whether these variants affect the binding and enzymatic activities of ResT during telomere resolution. Further studies are required to address this question.

### Plasmid DNA insertion at the right end of the chromosome: A new insight into Canadian strains

This study provided important new information about the *Bb* chromosome of Canadian strains: the insertion of a plasmid-like sequence at the right telomeric end of the linear chromosome. Although we observed that the linear chromosome across the nine strains is relatively stable, the most striking difference is observed at the right end, with genetic rearrangements between ∼865 kb to 877 kb (Figure 4B) and a plasmid-like DNA insertion at 900 kb (Figure 5). We identified that the telomeric sequences at the extreme right end are 99% identical to the terminal DNA segments of linear plasmid lp28-1 from strain 64b. This type of plasmid-like DNA segments are also found in other *Bb* strains. For example, right telomeric end of the reference strain B31 is >98% identical to the plasmid lp28-1 of strain 297, whereas the strain 297 carries a right telomeric end >99% identical to the B31 plasmid lp21 (Casjens et al. 2012). In contrast, the reference strains N40 does not carry any plasmid-like DNA extension, indicating that this feature is not universal. Interestingly, we observed similar variations in the plasmid-like DNA insertion among the assembled strains where four strains (Bb16-131, 133, 134 and 33) carried a much shorter DNA extension compared to the others. Although the precise mechanism of joining such plasmid sequences to the chromosome end remains unknown, and why it seems restricted to right end, Huang et al. ((Huang 2004)) suggested that it might result from recombination between the chromosome’s right end and linear plasmids. The high similarity (>99%) between the exchanged DNA segments, their proximity to the end, and the intact nature of the transferred genes suggest a homologous recombination event rather that non-homologous, which would be unlikely to maintain complete gene integrity (Huang 2004; Casjens et al. 2012; Casjens 1999). However, several questions remain unanswered. Does the increased genomic plasticity at the right chromosome end benefits *B. burgdorferi*? How does incorporating parts of the lp28-1 plasmid influences the pathogenicity, especially since some strains carry more integrated genes than the others? The answer probably lies in what we know about the function genes integrated from lp28-1. A previous study reported that the loss of the lp28-1 plasmid in strain B31 is correlated to decrease infectivity (Labandeira-Rey and Skare 2001). In this regard, the 2-10 kb DNA segment integrated into the chromosomes does not carry the surface antigenic variant generation system (*vlsE*), which is crucial for evading the host immune response; however, the integrated genes may still contribute to the overall infectivity.

### Unveiling a novel plasmid subtype

Despite detecting the integration of part of the lp28-1 plasmid into the chromosome of the nine strains, we were unable to assemble a complete lp28-1 plasmid for these strains (Table 2). This absence could be due to the plasmid being lost during the culturing process or decaying after contributing a substantial portion to the chromosome. While the precise mechanism remains unclear, our identification of a novel lp28-1a subtype plasmid in Bb16-22 offers some insight. In this novel plasmid, approximately 35% of the total length originates from lp28-1 and includes the *vlsE* system, while the remainder consists of sequences from five other linear plasmids (Figure 7D). This type of fusion among linear replicons was previously described for *Borrelia*, where plasmids often share highly similar sequences with others while their immediately adjacent sequences differ significantly (Casjens et al. 2012). What is particularly novel about our observation is that, while a part of lp28-1 is integrated into the chromosome, part the of the remainder sequences can be fused along with segments from other linear replicons to form a unique plasmid subtype. Some questions arising form these observations are whether this new lp28-1a subtype plasmid would be stable during *in-vitro* propagation and whether this new plasmid subtype is restricted to particular geographical regions. Additionally, it is tempting to speculate that the presence of this plasmid may contribute to the infectivity of *Borrelia* Bb16-22, given that it carries the *vlsE* gene cassette from lp28-1. Future studies will be required to investigate the biological implications of this novel plasmid subtype.

### In-depth comparative analyses are necessary to reveal the hidden genomic diversity

Comprehensive comparative genomics analyses are essential for uncovering the true genetic diversity within *Borrelia* strains. In this study, we demonstrate how telomere-to-telomere complete genome assemblies can enhance our ability to characterize genomic variations among *Borrelia burgdorferi* strains. Given that the nine assembled strains were collected from geographically proximate locations, we initially expected their genomes, particularly the linear chromosome, to be largely similar due to the typically conserved nature of *Borrelia* chromosomes. However, SNP-based phylogenetic analysis revealed unexpected variations even within the core conserved regions of the linear chromosome, indicating that strain relationships do not always strictly correlate with geographic origin.

Expanding our phylogenetic analysis to include additional Canadian *Borrelia* strains and incorporating both conserved and variable regions, we observed that the three Manitoba strains clustered together with previously assembled Manitoba strains, whereas the Ontario strains showed higher diversity with close relationships to the USA/Europe strains. Interestingly, one Ontario strain, Bb16-33, exhibited the greatest divergence from the other Ontario strains. These results suggest that genomic variation in the variable regions may be influenced by geographic factors.

To pinpoint the underlying sources of these variations, we conducted a detailed examination of multiple genomic features, including telomere length, conserved telomeric sequences, plasmid-like DNA segment extensions, plasmid profiles, and the diversity of key surface protein genes (*ospA*, *ospB*, *ospC*). These analyses revealed that, despite their relative geographic proximity, the nine assembled strains exhibit significant genomic diversity.

Overall, we developed a streamlined assembly pipeline that significantly reduces manual intervention while achieving high-quality, telomere-to-telomere genome assemblies of *Borrelia burgdorferi*, including comprehensive circular and linear plasmid assembly. By applying this pipeline, we generated nine complete *Borrelia* genomes from Canadian strains, providing a valuable resource for the scientific community studying Lyme disease in Canada and globally. Notably, our results reveal substantial genetic diversity among strains collected from relatively close geographical locations, underscoring the complexity of *Borrelia*’s genomic architecture. Based on the comprehensiveness of the assemblies, we recommend using Bb16-22 (Supplementary Figure S6) as a reference strain for future studies on Canadian *Borrelia*, especially those focusing on genomic diversity and pathogenicity. These findings highlight the importance of using complete genome assemblies to uncover the genetic diversity within *Borrelia* populations and provide a foundation for future research to understand their evolutionary dynamics and implications for disease transmission and virulence.

## Acknowledgements

We thank the Public Health Agency of Canada for generously providing the Bb strains and the McGill Genome Center for supporting with the sequencing facilities. We thank Aida Minguez Menéndez for her expert support in figure preparation. Atia Amin is supported by a Vanier Canada Graduate Scholarship. This work was funded by the McGill Interdisciplinary Initiative in Infection and Immunity (MI4) program grant to MN, DL, CFP and MO, the Foundation of the MUHC, the Louise and Alan Edwards and R. Howard Webster Foundation, which also supports MN. DL is also supported by a Fonds de la Recherche Québec-Santé (FRQ-S) Chercheur-Boursier Junior 2 Award and the Canada Foundation for Innovation John R. Evans Leaders Fund.

## Author Contributions

AA, CFP and DL designed the study, prepared the first draft of the manuscript, and revised the final draft of the manuscript. AA, AVIM, SG, GM and DL performed the experiments and analyzed the data. MO and MN contributed to the design, analyses, and manuscript revision. CFP, MB, and DL supervised the work. All authors have read and approved the content of this manuscript.

## Supplementary Figures

**Figure S1.**
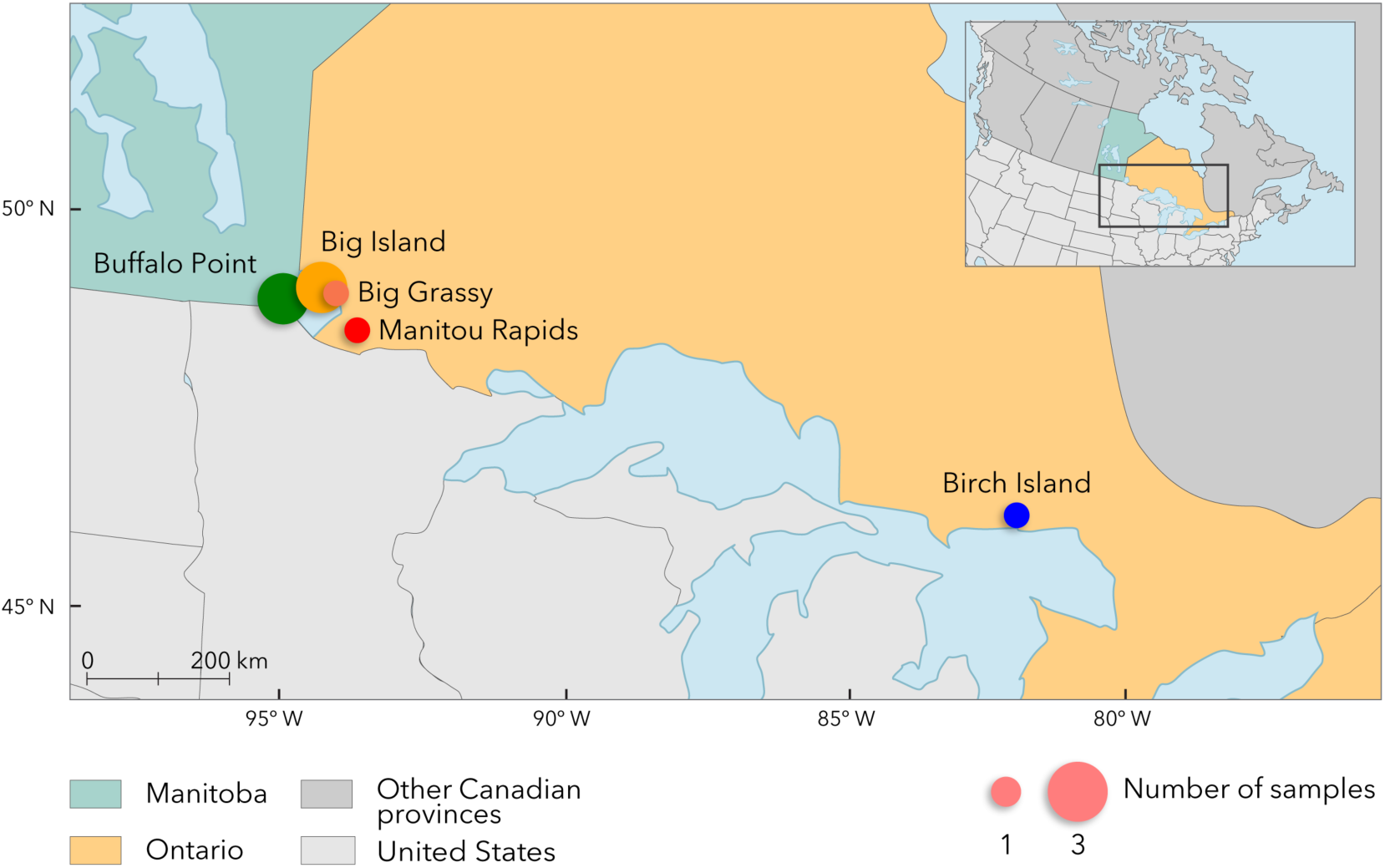
Geographical origins of the collected *B. burgdorferi* samples. Three samples were collected from Manitoba and six from various locations in Ontario. Distinct colors are used to represent different sample collection sites.

**Figure S2.**
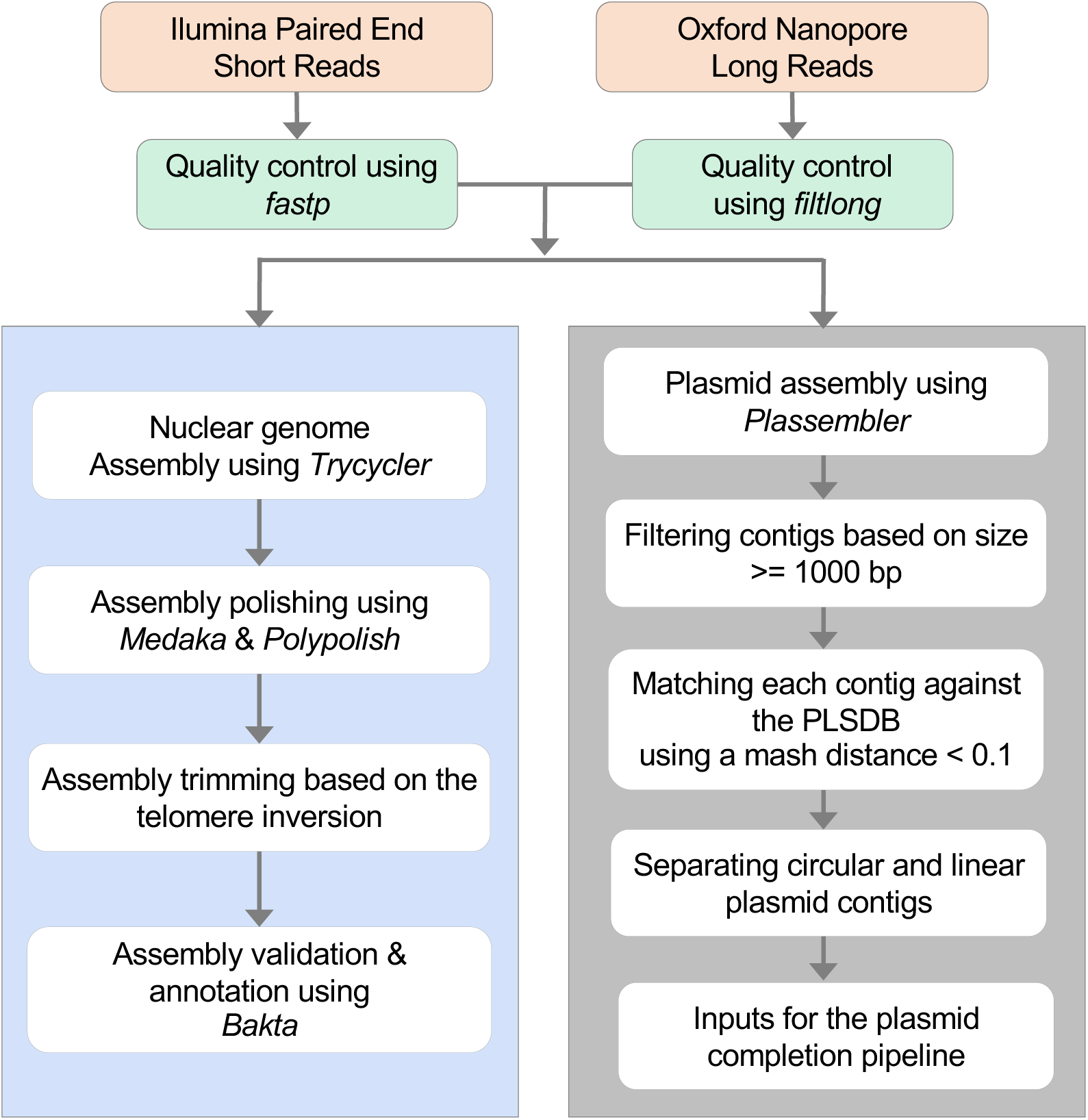
Genome assembly pipeline developed in this study. The orange boxes refer to the input files, while the green boxes refer to the tools used for quality control of the raw sequencing data before incorporating them into the assembly pipeline. The blue and grey boxes represent the linear chromosome assembly and plasmid assembly pipelines, respectively. The names of the tools are italicized to distinguish them from the files.

**Figure S3.**
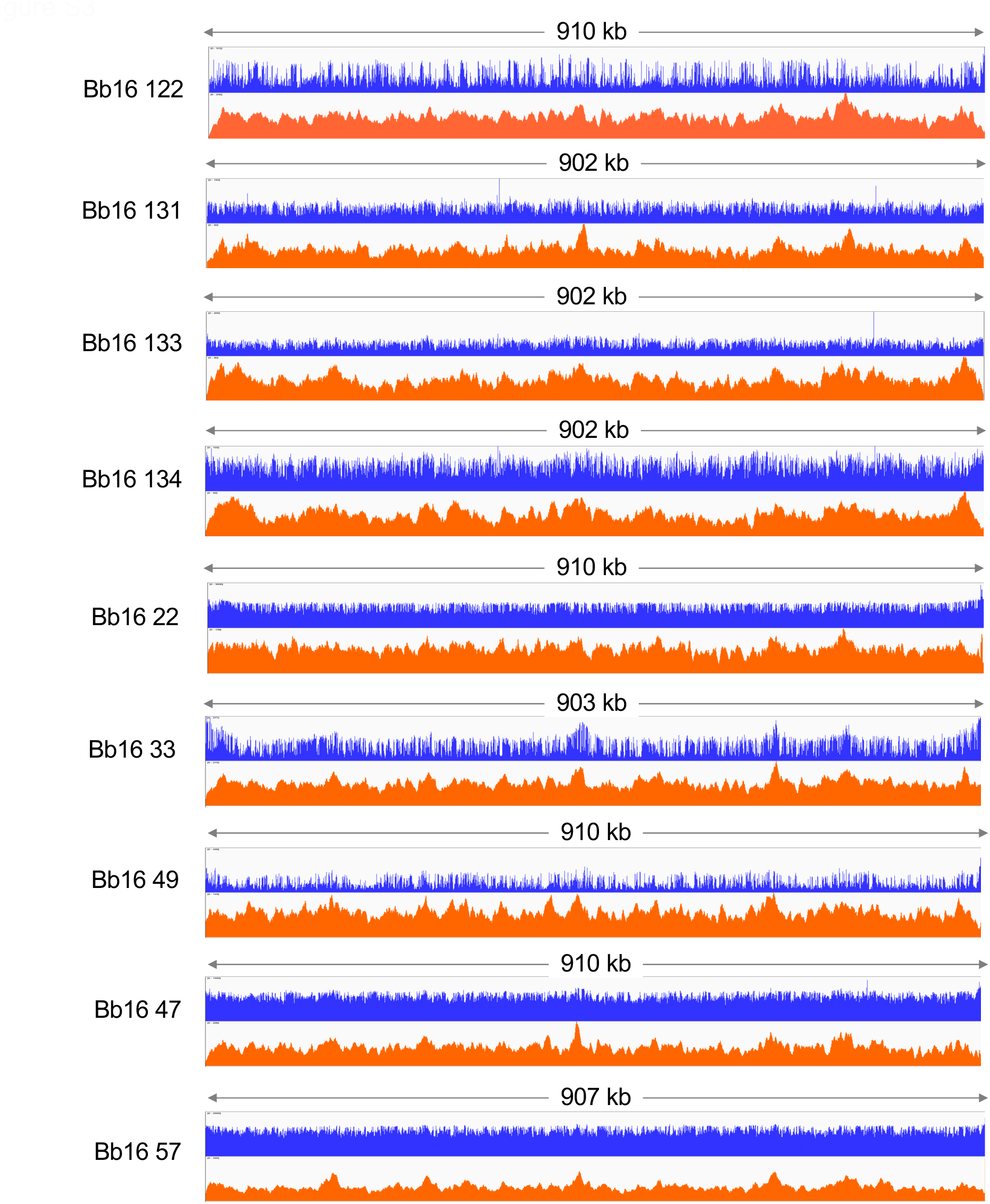
Read coverage across assembled linear chromosomes. Illumina (blue) and Oxford Nanopore (orange) sequencing read coverage across the fully assembled linear chromosomes of the nine Canadian strains.

**Figure S4.**
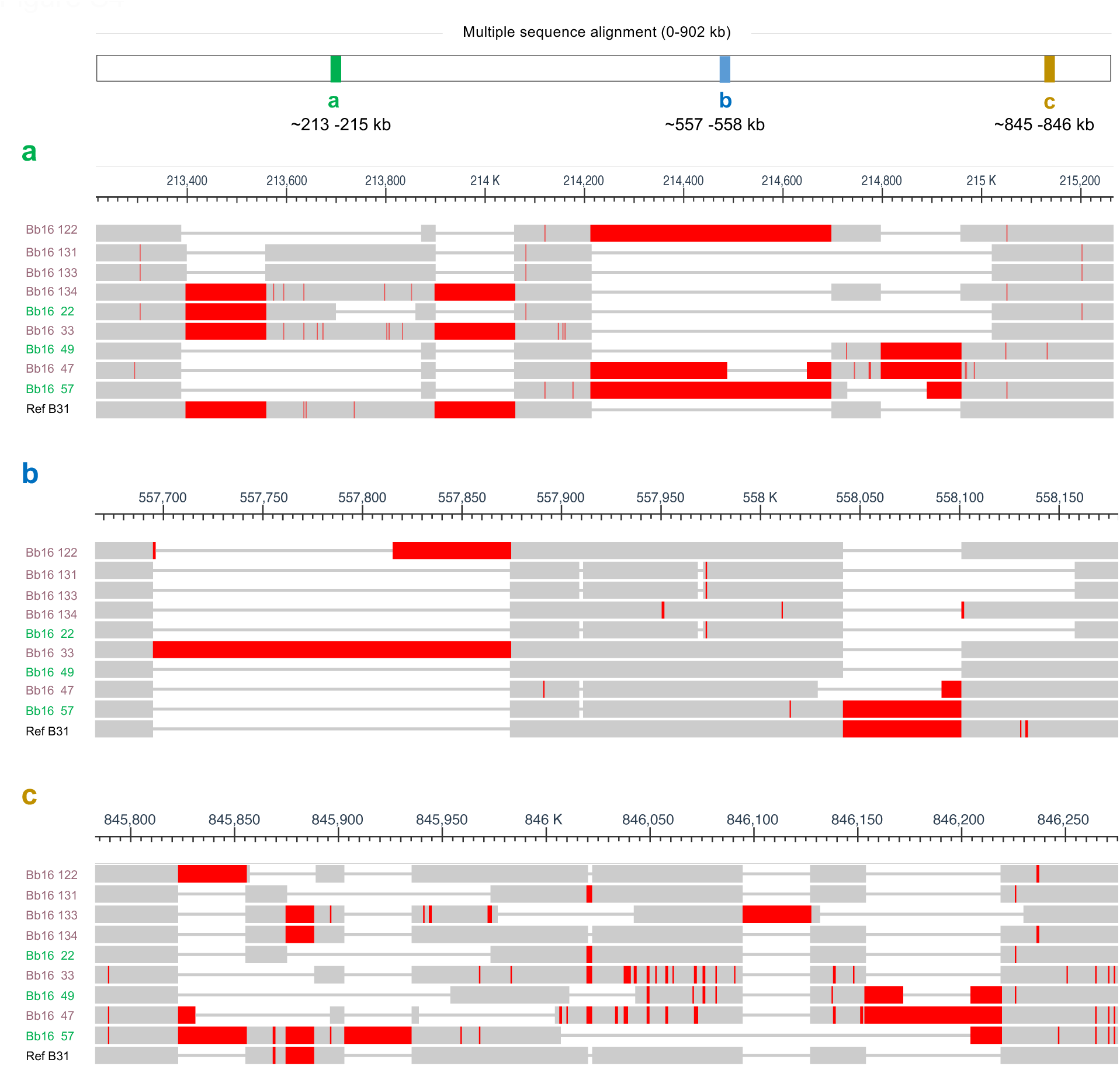
Detection of variable regions within the conserved linear chromosome. Three variable regions (a, b, and c) were identified within the conserved regions of the linear chromosome, using a comparative analysis between the nine assembled strains and the reference strain B31. Strain names are color-coded by geographic origin (green = Manitoba; pink = Ontario).

**Figure S5.**
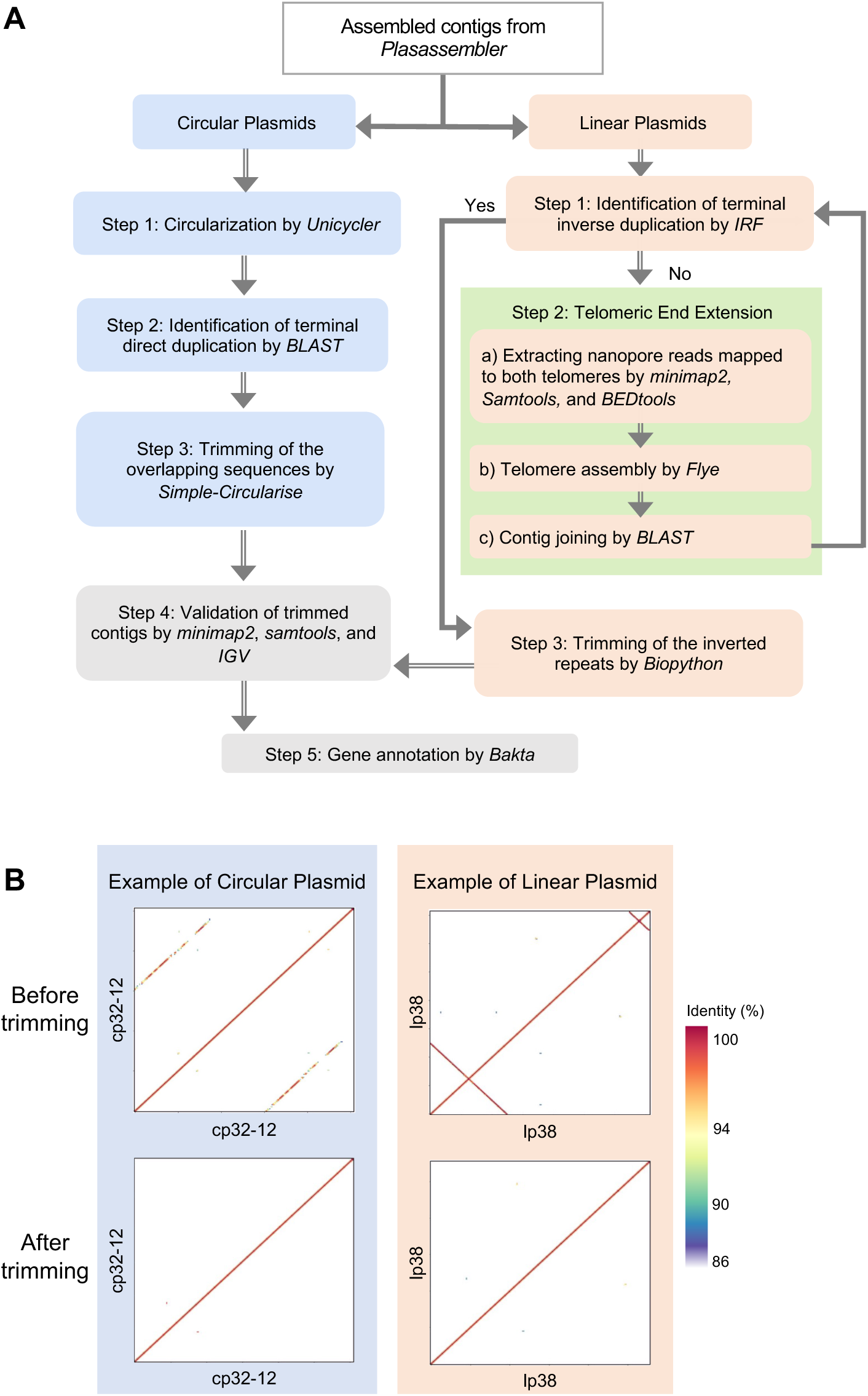
Trimming and circularization workflow for linear and circular plasmids. **A)** Schematic overview of the plasmid refinement pipeline, divided into separate steps for linear and circular plasmids. Blue boxes indicate steps specific to circular plasmids, orange boxes to linear plasmids, and grey boxes represent steps common to both. The green box denotes the “Telomeric end extension” step, which is unique to linear plasmids. Tool names are italicized. **B)** Examples of contig trimming for circular and linear plasmids visualized using self-identity dot plots generated with ModDotplot.

**Figure S6.**
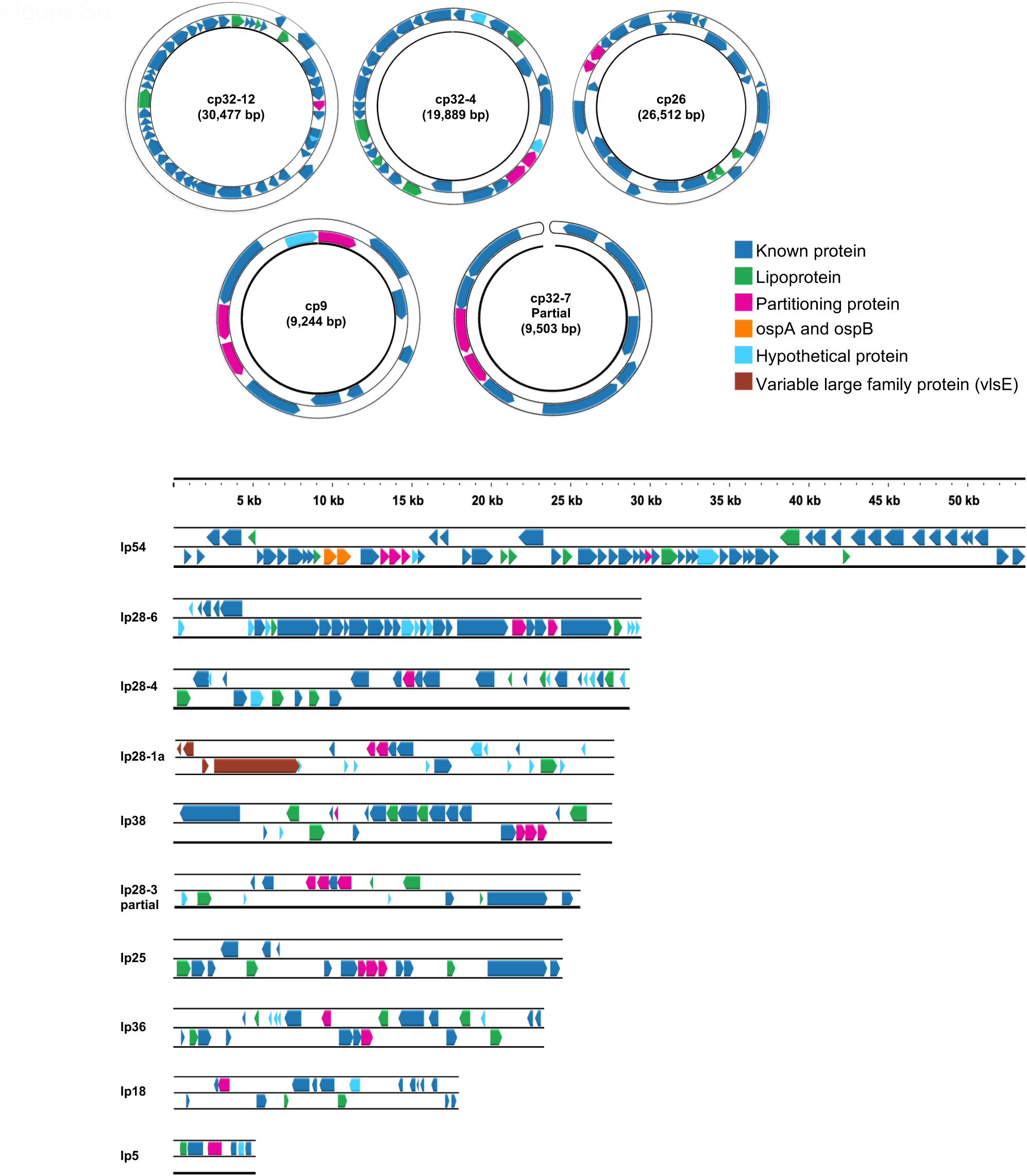
Visualization of the complete plasmid assembly and gene annotation of strain Bb16-22.

